# A structural model of the human serotonin transporter in an outward-occluded state

**DOI:** 10.1101/637009

**Authors:** Eva Hellsberg, Gerhard F. Ecker, Anna Stary-Weinzinger, Lucy R. Forrest

**Author notes:** Corresponding author (NINDS, NIH).

## Abstract

The human serotonin transporter hSERT facilitates the reuptake of its endogenous substrate serotonin from the synaptic cleft into presynaptic neurons after signaling. Reuptake regulates the availability of this neurotransmitter and therefore hSERT plays an important role in balancing human mood conditions. In 2016, the first 3D structures of this membrane transporter were reported in an inhibitor-bound, outward-open conformation. These structures revealed valuable information about interactions of hSERT with antidepressant drugs. Nevertheless, the question remains how serotonin facilitates the specific conformational changes that open and close pathways from the synapse and to the cytoplasm as required for transport. Here, we present a serotonin-bound homology model of hSERT in an outward-occluded state, a key intermediate in the physiological cycle, in which the interactions with the substrate are likely to be optimal. Our approach uses two template structures and includes careful refinement and comprehensive computational validation. According to microsecond-long molecular dynamics simulations, this model exhibits interactions between the gating residues in the extracellular pathway, and these interactions differ from those in an outward-open conformation of hSERT bound to serotonin. Moreover, we predict several features of this state by monitoring the intracellular gating residues, the extent of hydration, and, most importantly, protein-ligand interactions in the central binding site. The results illustrate common and distinct characteristics of these two transporter states and provide a starting point for future investigations of the transport mechanism in hSERT.

## Introduction

The serotonin transporter hSERT belongs to the secondary active solute carrier 6 (SLC6) membrane protein family, in which it forms the subgroup of monoamine transporters (MAT) together with the dopamine and norepinephrine transporters, DAT and NET [1,2]. The SLC6 transporters are human proteins belonging to the larger family of neurotransmitter:sodium symporters (NSS; Transporter Classification Database [3] identifier 2.A.22). The solute transported by hSERT is serotonin (5-hydroxytryptamine, 5HT), an important tissue hormone in the periphery and a neurotransmitter in the central nervous system. In its neurotransmitter function, 5HT plays a crucial role in regulation of, or impact on, mood, sleep-wake cycle, appetite, pain, sexuality, and body temperature control. hSERT is the primary target for competitive inhibitors in major depression therapy, but it also interacts with inhibiting, or even transport-reverting, psychostimulants. According to the WHO, depression is globally the leading cause of disability [4]. Despite its importance, the serotonin transport mechanism is not yet fully understood. 5HT is released from vesicles in presynaptic neurons into the synaptic cleft, where it transmits its signal to the postsynaptic 5HT-receptors. After transmission, 5HT is taken back into presynaptic neurons for degradation or vesicle storage. This reuptake against its concentration gradient is facilitated by hSERT under cotransport of sodium [5,6]. Furthermore, chloride is required for transport activity [7], while potassium antiport stimulates the transport process [8]. However, the exact transport stoichiometry remains elusive and the potential binding site for potassium in the transporter is unknown; both questions need to be addressed for a complete understanding of the transport mechanism. To facilitate 5HT reuptake, the transporter needs to undergo distinct conformational changes. In principle, these changes expose the substrate binding site(s) to one side of the membrane at a time, according to the so-called alternating-access mechanism [9], support for which has been provided by structural modeling and biochemical experiments [10–12], and recently also by X-ray and cryo-EM crystallography [13–15]. Taken together, the hSERT transport cycle can be represented roughly as shown in **Fig 1**, albeit with the caveat that several important details remain elusive.

**Fig 1:**
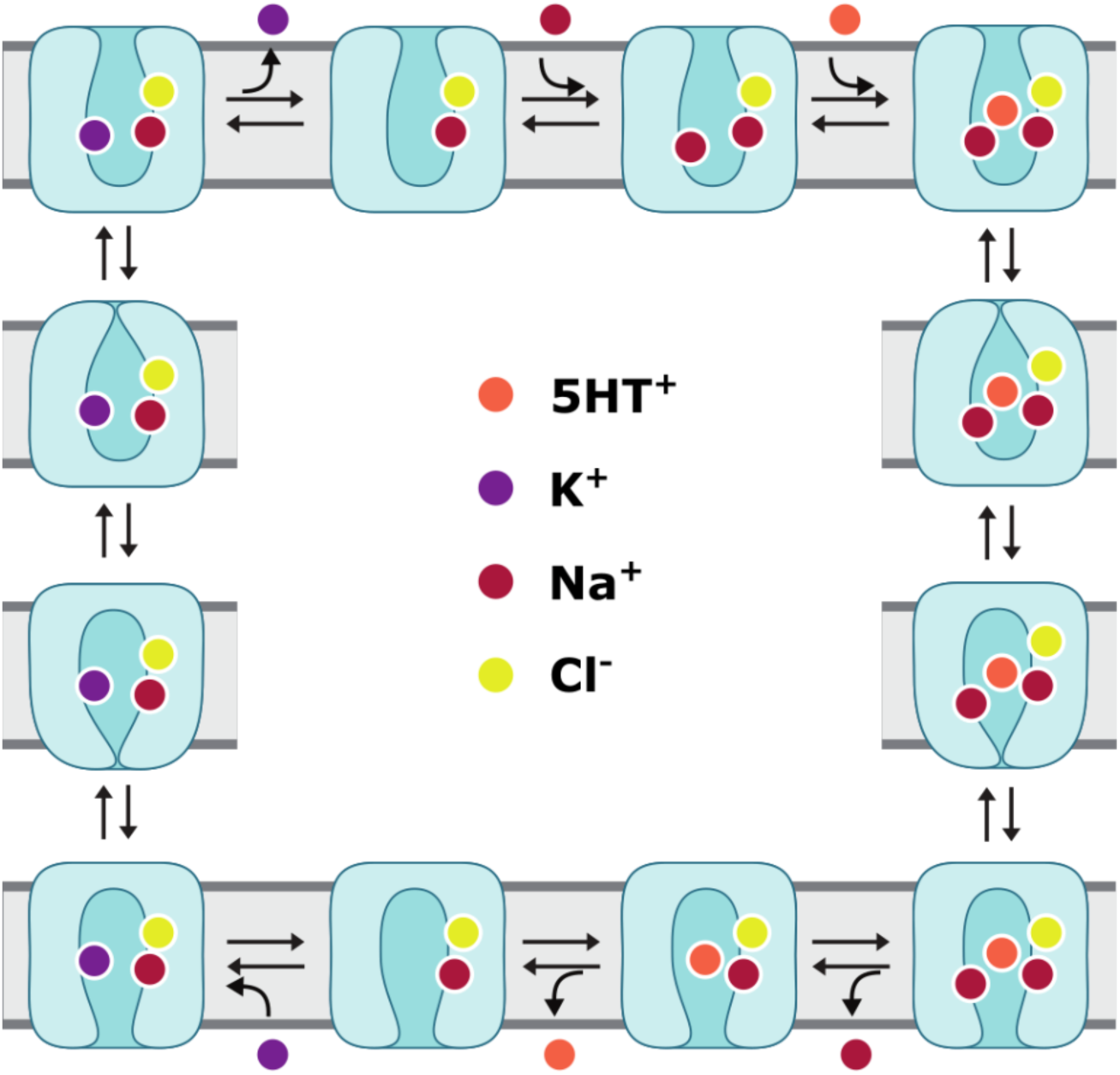
Schematic representation of key steps in the physiological transport cycle of hSERT. The outward-open conformation of hSERT bound to Na^+^ and Cl^-^ is entered by a second Na^+^ ion and the endogenous substrate 5HT^+^. Once all these required components are bound, the extracellular gates can close, allowing the transporter to undergo progressive conformational changes through the outward-occluded, fully-occluded, and inward-occluded transporter states. Once the inward-open state is reached, one Na^+^ ion and 5HT^+^ are released into the cytosol. Upon binding of a K^+^ ion, the transporter resets via equivalent conformational changes, but in reverse order, finishing its cycle by release of K^+^ into the synaptic cleft.

hSERT belongs to a group of transporters sharing the LeuT structural fold [16], named for the eubacterial *Aquifex aeolicus* amino acid transporter LeuT_Aa_ [17]. The LeuT fold comprises ten transmembrane helices (TMs), that contain a structural repeat involving TM1-TM5 and TM6-TM10, which are oppositely oriented in the membrane (**Fig 2**). This architecture has also been divided further into the so-called scaffold and bundle domains, according to the proposed “rocking bundle” mechanism [18]. However, it should be noted, that different LeuT-fold transporters exhibit variations in the domains involved to facilitate translocation [19].

**Fig 2:**
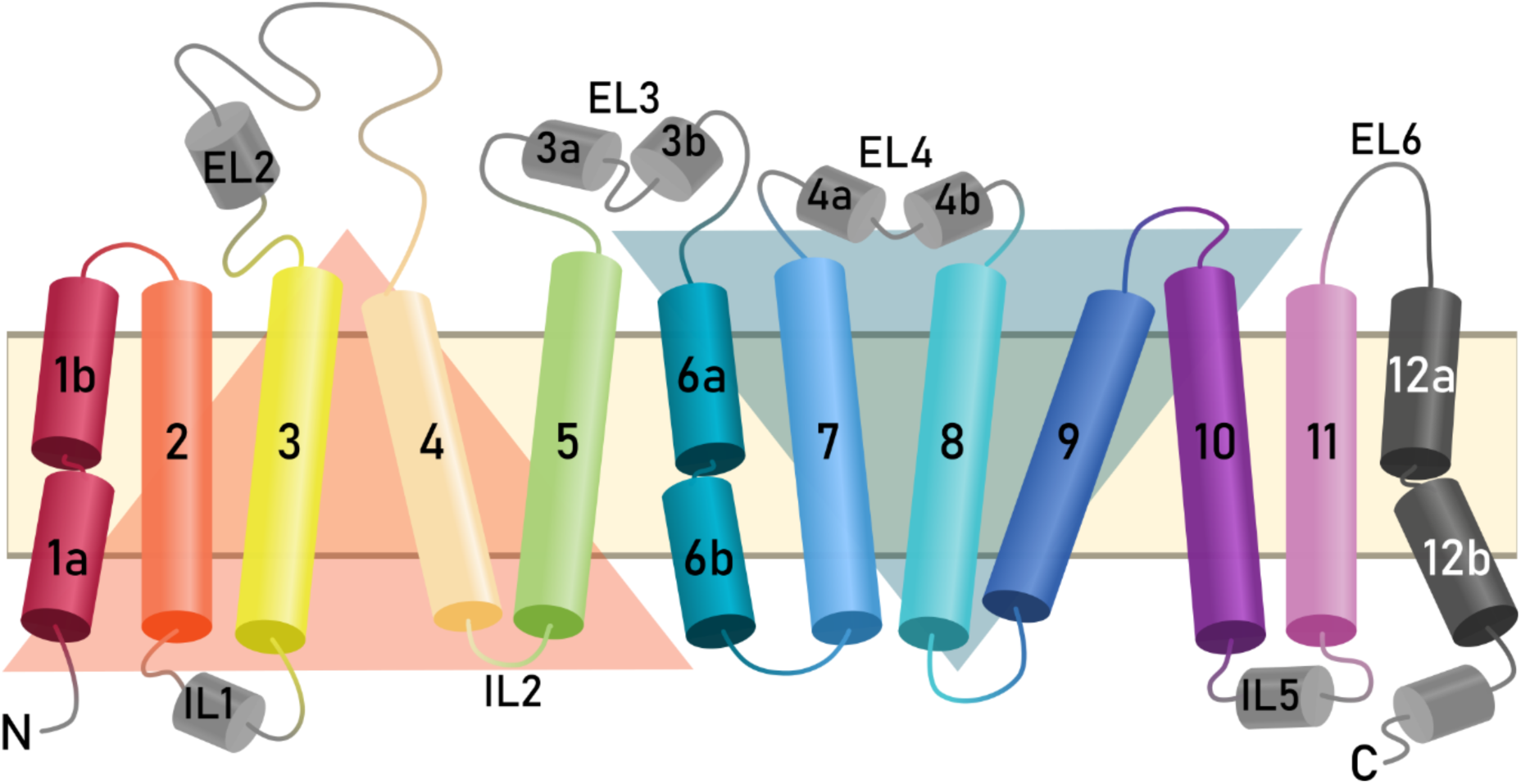
Transmembrane topology of hSERT comprising twelve membrane-spanning helices connected by intra- and extracellular loops (ILs, ELs). The inverted-topology structural repeats are highlighted by orange (TM1-5) and blue (TM6-10) triangles. The “rocking bundle” proposed to facilitate substrate transport consists of TM1, TM2, TM6, and TM7.

In hSERT, the bundle helices, comprising TM1, TM2, TM6, and TM7, are proposed to undergo the required conformational changes to facilitate the transport of 5HT, during which they form and break molecular interactions according to a specific gating mechanism. The gating residues are located in the intra- and extracellular vestibules of hSERT and connect or disconnect the bundle to or from the scaffold [20]; these residues are also highly conserved across the NSS family [21]. These gates were identified by mutational studies, including binding affinity and solvent accessibility measurements in hSERT [10,18,22,23] and were observed in structures of LeuT_Aa_ and *Drosophila melanogaster* DAT in different conformations [17,24–38], prior to the release of the hSERT structures. In the extracellular vestibule, there are two gates: R104 (TM1b) and E493 (TM10), which can form a salt bridge, and Y176 (TM3) and F335 (TM6a), which can form a hydrophobic lid directly covering the bound substrate from the extracellular side. In the intracellular pathway leading to the cytoplasm, there is a comprehensive network of residues involving parts of TM5, TM6, TM8, TM9, and the N-terminus. The interaction pairs responsible for opening and closing of the cytoplasmic permeation pathway feature the most conserved residues, namely R79-D452, W82-Y350, Y350-E444, and E444-R462.

Since the structures of LeuT_Aa_ and dDAT were reported, many experimental and homology-based computational studies have been carried out to explore the transport mechanism, stoichiometry, and ligand recognition patterns of hSERT [10,12,39–59]. Nevertheless, important questions remain unresolved, including the structure of the substrate-bound transporter in key conformational states (as opposed to inhibitor-bound conformations); the conformation of 5HT in its orthosteric binding site; and the interactions required for the substrate to facilitate coupling and conformational change. To address these questions, we built a comparative model of hSERT in an outward-occluded state in complex with 5HT. To this end, we applied a procedure involving two different template structures [56]. Comparison of the behavior of the resultant model during molecular dynamics simulations with that of the outward-open conformation also bound to 5HT, allows us to predict the structural features that characterize this state and that differentiate it from the outward-open state. The very recent cryo-EM data showing hSERT in an outward-occluded conformation bound to the inhibitor ibogaine provides support for many features of the model and suggests subtle ligand-dependent differences in this conformation.

## Methods

### Structural modeling

To build a model of hSERT (Uniprot identifier P31645, SC6A4_HUMAN [60]) in an outward-occluded state, we combined the available information from the outward-open X-ray structure of hSERT (PDB entry 5I71 chain A, at 3.15 Å resolution, referred to hereafter as 5I71) and the outward-occluded X-ray structure of LeuT_Aa_ (PDB entry 3F48 chain A at 1.9 Å resolution, referred to hereafter as 3F48). As explained in the introduction, the extracellular half of the bundle is responsible for the conformational change from an outward-open to an outward-occluded state. Therefore, helices TM1b and TM6a, as well as the extracellular halves of TM2 and TM7 were extracted from the outward-open hSERT, and repositioned based on the positions of the corresponding regions of the outward-occluded LeuT_Aa_ structure. EL3 and EL4a were repositioned in the same manner, to maintain their helical packing against other helices in the bundle. All remaining protein segments, namely TM1a, IL1, EL2, TM3, TM4, TM5, TM6b, EL4b, TM8, TM9, TM10, IL5, TM11, TM12, the intracellular halves of TM2 and TM7, as well as the N- and C-termini were taken directly from the outward-open structure of hSERT (**Fig 3**). To assign the boundaries of the extracellular bundle components, hSERT 5I71 and LeuT_Aa_ 3F48 were first aligned using the backbone atoms of the hash region, which comprises TM3, TM4, TM8, and TM9 [61], in VMD v1.9.3 [62]. The boundaries were assigned by taking into account sequence similarity and the preservation of secondary structure.

**Fig 3:**
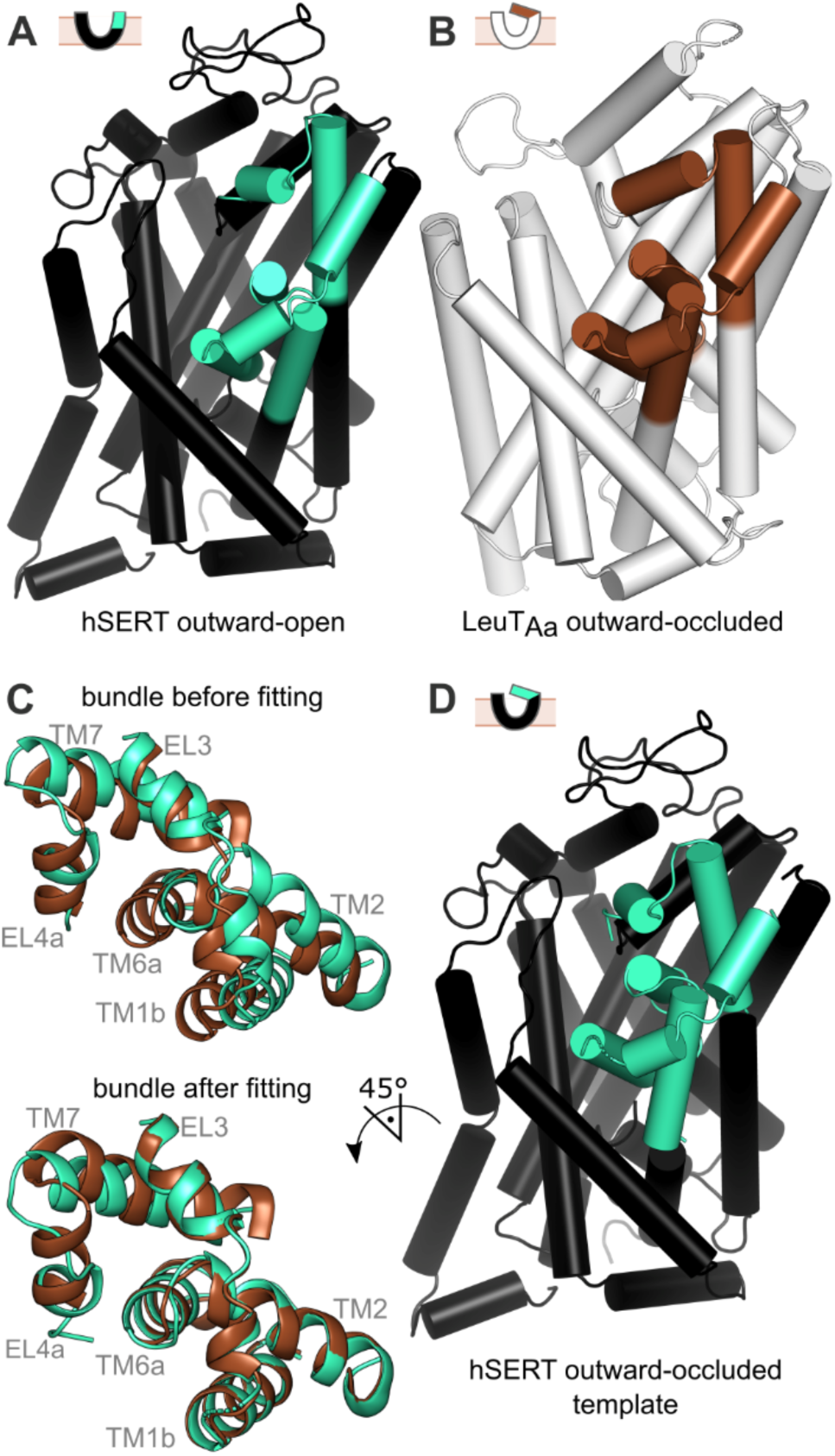
Model-building process. (A, B) The two template structures used were outward-open hSERT (A, PDB ID: 5I71) and outward-occluded LeuT_Aa_ (B, PDB ID: 3F48). The extracellular bundle components whose conformations define the occlusion of the extracellular pathway in NSS transporters are shown in green (hSERT) and brown (LeuT_Aa_). C) To reposition the outward-open hSERT bundle segments according to their expected locations in the outward-occluded state, the former were individually superposed onto the corresponding elements of LeuT by structural alignment. D) The resultant modeling template consists of the unchanged segments (black) plus the now outward-occluded, repositioned bundle segments (cyan), both originate from hSERT itself.

After testing different combinations of segments, the best fit – defined as the smallest root mean squared difference (RMSD) for the largest combination of fragments – between the extracellular halves of the bundle in hSERT and the equivalent segments in LeuT_Aa_ was achieved by dividing the bundle and the extracellular loops into five segments (S1 Table). Modeller v9.15 [63] was then used to build the outward-occluded hSERT model, by combining the repositioned outward-occluded extracellular bundle segments of hSERT with the remaining components of hSERT 5I71, and optimizing their physicochemical properties. The input alignment for Modeller derived from the template building process can be found in S2 File. A total of 2000 models were generated, sufficient to reach convergence of the Modeller molpdf score. The molpdf score was also used for filtering out a subset of models for further analysis. The ProQM score [64] predicts the quality of individual models with both local (residue) and global (protein) resolution and was used for further model selection. Lastly, Ramachandran plots generated with PROCHECK [65] were used to assess the protein backbone quality; models with residues in disallowed regions were excluded from further analysis.

### Model refinement and final selection

To prepare the selected models for docking and simulation, we added ions into the Na1, Na2 and Cl binding sites and refined the surrounding binding-site regions accordingly. The sodium and chloride ions were transferred from the available crystal structures by aligning the respective binding sites including all residues within 4.5 Å onto those in the models. The chloride ion was taken from the hSERT PDB entry 5I6X (chain A, referred to hereafter as 5I6X), and the two sodium ions were taken from hSERT 5I71. From the latter, a water molecule within H-bonding distance of D437 was also copied. The side chain rotamers of the ion-coordinating residues were set to those in the X-ray structures by adjusting the corresponding dihedral angles. Similarly, the side chain orientations of F334 and F335 were changed to those of F252 and F253 in LeuT_Aa_ 3F48 to reflect the outward-occluded conformation. The native residue T439 was reintroduced instead of the mutation S439.

The models were energy minimized using 1000 steps of steepest descents in three iterations including first only hydrogen atoms, then side chain atoms, and lastly, all atoms. The distances between the coordinating atoms of the binding sites and the ions were restrained to the values obtained from the corresponding X-ray structures with a force constant of 10 kcal/mol/Å^2^ (S3 File). Side chain rotation, mutation, and energy minimization were conducted in CHARMM 36b2 [66] because of its accuracy in placing hydrogens [67]. Finally, the models were analyzed with MolProbity v4.4 [68,69] to check for steric clashes, expected side chain orientations, and proper ion coordination.

### Induced-fit docking (IFD) of 5HT into outward-occluded models

5HT was placed into the orthosteric binding site by molecular docking. This binding site was defined as including all residues within 4.5 Å of paroxetine in hSERT 5I6X (Y95, A96, D98, A169, I172, A173, Y176, F335, S336, L337, G338, F341, S438, T439, G442, T497, V501) using Maestro [70]. No further protein preparation was performed as the structures were already protonated and energy-minimized. The protonation state of 5HT was predicted using the chemicalize web service [71]. With this method, the primary amine of 5HT was predicted to be protonated 99.5% of the time at pH 7. The structure was prepared in LigPrep [72] with default options. The induced-fit docking [73] was carried out for multiple protein models with the extended sampling protocol, which generates up to 80 poses, using automated docking settings. We used the OPLS3 force field and kept the default options, i.e., no constraints, ligand conformational sampling within 2.5 kcal/mol, a softened van der Waals potential (radii scaling), Prime refinement of residues within 5 Å of the ligand, and Glide redocking of all poses within 30 kcal/mol of the lowest-energy pose. All poses obtained for all models were clustered pair-wise according to the non-hydrogen atoms of 5HT and the backbone atoms of the binding site residues using the conformer clustering module of Maestro [70]. Thus, the clusters were generated with the average linkage method without fitting, and the cluster number was determined at a minimum of the Kelley function [74]. The resultant clusters were analyzed according to cluster size, docking scores, and orientation of 5HT according to the findings of Celik et al. [45] using interaction fingerprints.

### Molecular dynamics (MD) simulation setup

Simulations were set up using the CHARMM-GUI web service [75] with a few adaptations described in this section. First, the protein structures were aligned relative to the membrane by using VMD to fit them to the structure of hSERT (5I6X) obtained from the OPM database [76], using only the backbone atoms of the hash region. In the CHARMM-GUI membrane builder [77], the C-terminus was acetylated and the N-terminus amidated (ACE and CT2 caps). The native residues Y110 and I291 were reintroduced instead of A110 and A291, respectively. Y110 became distorted during membrane embedding and was rebuilt with the Swiss-PdbViewer v4.1 [78]. E508 was set to the protonated form, as it was reported to H-bond with E136 [79]. C200 and C209 were connected with a disulfide bond. We used a phosphatidylcholine lipid bilayer (POPC), solvated with TIP3P water and 0.15 M NaCl. Half of the bulk sodium ions were replaced with potassium ions, reflecting the ability of hSERT to countertransport potassium. Transporter cavities were solvated with Dowser [80]. The final system contained ~140,000 atoms including 330 lipids, 38 sodium, 39 potassium and 83 chloride ions as well as 28,985 water molecules, with box dimensions of ~11 nm^3^ after equilibration (see below). Ligand parameters for 5HT (S4 File) were generated with the CHARMM general force field (CGenFF) v3.0.1 [81–83] using the ParamChem web server v1.0.0.

### Molecular dynamics (MD) simulations

Following the default CHARMM-GUI protocol for Gromacs with the CHARMM36m force field [84], steepest descents energy minimization and six equilibration steps comprising gradual reduction of side chain and backbone restraints in steps 1-3 for 25 ps and steps 4-6 for 100 ps each, were performed. Subsequently, two additional equilibration steps were carried out using the PLUMED v2.3 plugin [85] to apply specific restraints with a force constant of 200 kcal/mol/Å^2^ over a simulation time of 500 ps each, as follows. During the first equilibration step, restraints were applied to relevant distances in the orthosteric and ion binding sites, as well as between the extracellular gate residues Y176 and F335, and R104 and E493 (S5 Table). The second equilibration step allowed the protein backbone to tumble in the membrane by the use of an RMSD restraint. Energy minimization, equilibrations, and production runs were carried out with Gromacs 2016.3 [86–88] using a 2 fs time step, and the trajectory output was saved every 20 ps for the analyses. In the production runs, a temperature of 301 K was maintained using velocity rescaling and pressure was controlled by semi-isotropic coupling using Parrinello-Rahman barostat; the Berendsen barostat was used during equilibration. A Verlet neighbor search was used for van der Waals and Particle Mesh Ewald electrostatics interactions within 1.2 nm. Van der Waals interactions were switched above 1.0 nm. Bond lengths were constrained with the LINCS algorithm.

For the production runs with the outward-occluded models, we carried out a number of test simulations to identify complexes in which the pathway remained closed (as tracked by the extracellular gate distances). We identified one 2-microsecond long trajectory in which the pathway remained closed throughout; production runs were initiated from four time-points during that trajectory (at t = 500, 600, 800 and 1000 ns, respectively) and were carried out for one microsecond each. For the outward-open complex (see below), we carried out four replica production calculations, each one microsecond in length, with differing initial velocities. Final frames from each trajectory are provided as supporting information S6 and S7 Data.

### Outward-open structure preparation, docking, and simulation setup

Overall, the methods used for docking and simulations of the hSERT outward-open crystal structure (5I71) were the same as for the outward-occluded models, with the following exceptions. After structure preparation in CHARMM-GUI, the structure was minimized with the Protein Preparation Wizard in Maestro [89,90] using default options. The simulation setups were prepared in the CHARMM-GUI web service [75] without further adaptations. Energy minimization, equilibrations and production runs were carried out with Gromacs 5.1.1.

### Analysis of water positions

Water molecules entering the protein interior were counted using an in-house tcl script for VMD v1.9.3 following the same method applied previously [91], except that the protein interior, defined as a 20 x 15 x 25 Å box, was divided into intra- and extracellular vestibules, each a length of 9 Å along the z-axis, and the orthosteric binding site, which was 7 Å long. The residues used to define the approximate boundaries of the orthosteric site were: Y176 in TM3 and F335 in TM6a on the extracellular side and F341 in the unwound region of TM6 on the cytoplasmic side. The extracellular vestibule was defined as reaching as far as R104 in TM1b and E493 in TM10, while the intracellular vestibule was defined as extending to L89 in TM1a and S277 in TM5.

Water occupancy was mapped using the VolMap tool in VMD v1.9.3, after aligning the protein using all backbone atoms. All water molecules within 20 Å of any atom in 5HT in the orthosteric binding site were included for map generation at a resolution of 2 Å and averaged over all frames in each trajectory.

## Results

### A structural model of hSERT in an outward-occluded state

We created a comparative model of hSERT in an outward-occluded conformation by combining information from outward-open structures of hSERT and outward-occluded structures of LeuT_Aa_ (**Fig 3**). An outward-open structure of hSERT bound to S-citalopram (5I71) was chosen, as the structure contains ions resolved at both Na1 and Na2 sites, which is important because these sites are located directly at the interface region where the conformational change is most substantial. For LeuT_Aa_, a structure of the outward-occluded conformation (3F48) was used, because it is bound to alanine, which has the highest transport turnover rate among the LeuT_Aa_ substrates. The majority of the template components were reproduced exactly as in the hSERT structure (5I71), namely the scaffold regions not expected to be involved in this conformational change, as well as the entire intracellular segment of the protein. The extracellular bundle of hSERT responsible for the occlusion of the extracellular pathway was superposed onto the corresponding region in LeuT_Aa_ such that the repositioned coordinates of this region of hSERT could be used as a template. The two templates were then connected in order to build the “homology” model of hSERT in an outward-occluded state.

Of the 2000 models built, the best 100 models according to the Modeller molpdf score, i.e., those that best satisfy all the input restraints, were selected for further analysis. Based on the local and global ProQM scores (mean 0.817, stdev. 0.005), these models are of comparable quality to the X-ray structures used (0.819 for 5I71 and 0.796 for 3F48). To filter the selection of models further, we selected the 16 models with scores within one standard deviation from the highest global ProQM score; these scores fall in the range from 0.822 to 0.827 (**Fig 4**). In all remaining models, 92-94% of their residues are found in the favored regions of the Ramachandran plot; out of these models, we selected the eight models with no residues in disallowed regions.

**Fig 4:**
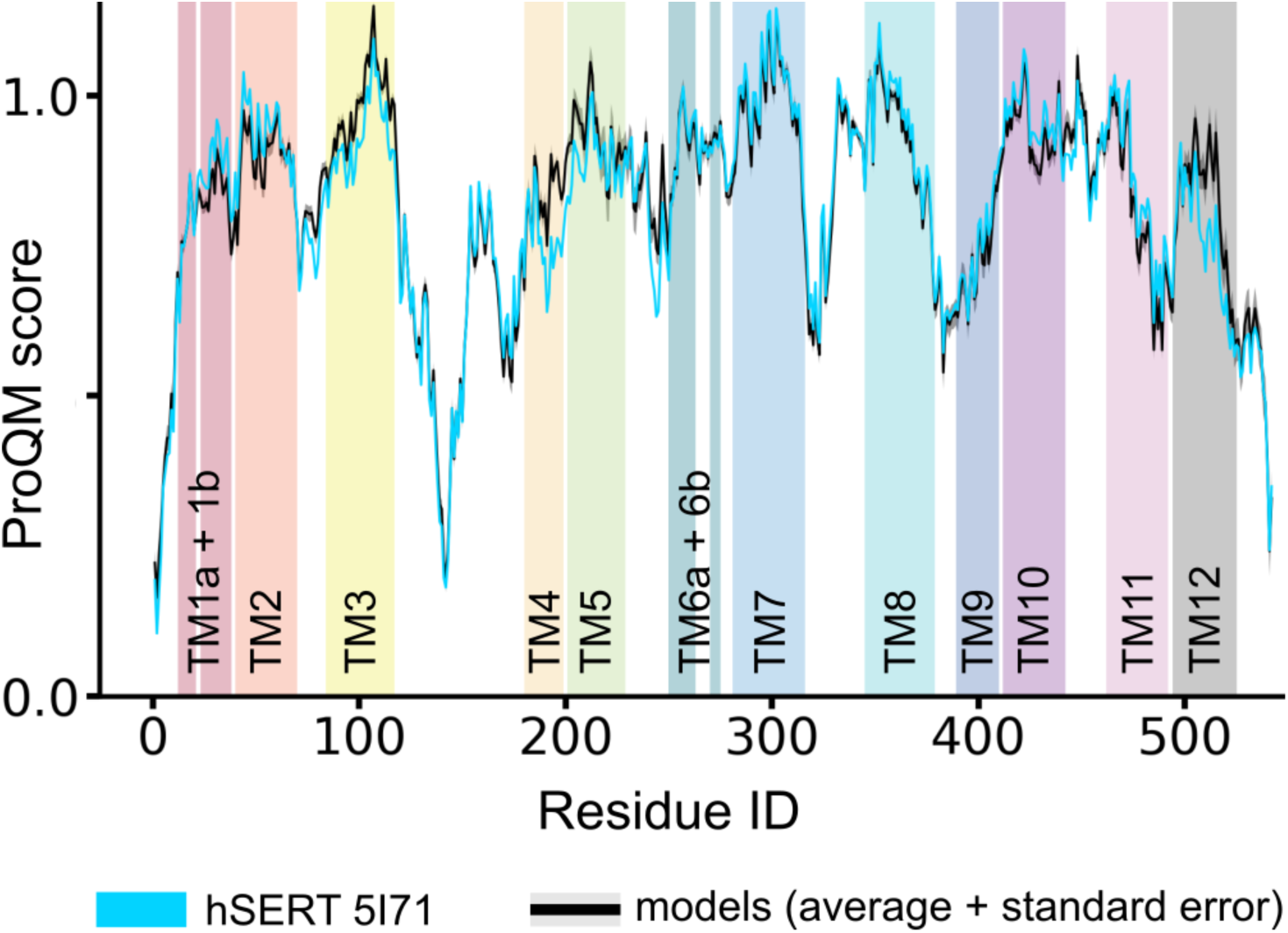
ProQM Plot. Plot of the local ProQM structural model quality scores for each residue in the X-ray structure (hSERT 5I71, cyan) and the 16 best ranked models (average in black, standard errors in gray). The ProQM plot shows a clear quality comparability between experimental and modeled structures and a striking consistency amongst the models.

After addition of sodium and chloride ions (in the Na1, Na2, and Cl sites), as well as a crystallographic water molecule, into all eight models, the side chains of binding-site residues (including F334 and F335) were rotated to match their orientations in known structures, and the resultant models were energy minimized. Energy-minimized models were assessed using MolProbity and selected for docking based on steric clashes, ion coordination distances, and the orientation of the F335 side chain rotamer. As Y176 and F335 form the extracellular lid of the orthosteric binding site, the orientation of F335 is a crucial characteristic for any transporter state that is closed to the extracellular side [27,33]. By these criteria, three final models were selected for further studies (S8 Tables).

### Binding pose of 5HT in outward-occluded state

The occluded conformation of hSERT bound to sodium and chloride is physiologically only likely to be observed in the presence of the endogenous substrate, 5HT. To identify the orientation of 5HT in the orthosteric binding site, we carried out induced-fit docking to the three selected outward-occluded models and the outward-open X-ray structure of hSERT. This procedure resulted in comparable numbers of poses, docking scores (Glide Gscores and IFD scores), and cluster distributions, for both conformations of the protein. In one of the models, F335 adopted a different rotamer orientation as a result of the docking process, thereby exposing the orthosteric binding site to the extracellular pathway. We therefore excluded this model and continued using the poses of the two remaining models.

For analysis of the holo outward-occluded models, the 138 different docked poses were clustered according to the orientation of 5HT. Two clusters, containing > 15% of all poses (> 20 poses) each, were retained for further analysis. The highest-populated cluster, comprising 57 poses, contains poses in both models, as well as the best Glide and IFD scores on average, although the scores are very similar between clusters (S9 Tables).

Notably, the interaction fingerprints in the most-populated cluster are consistent with previous findings [45]. These interactions include: an ionic interaction (or with additional H-bond character) between D98 and the cationic nitrogen of 5HT; the C6 atom of 5HT positioned close to A173; an interaction between the 5-OH group of 5HT and T439; and the nitrogen atom in the indole ring of 5HT pointing towards F341. From the models in the best cluster, we selected the top-ranked pose for each of the two homology models as that with the lowest Glide Gscore (–9.1 and –8.9 kcal/mol, respectively) to be used as starting structures for subsequent MD simulations (**Fig 5A**).

**Fig 5:**
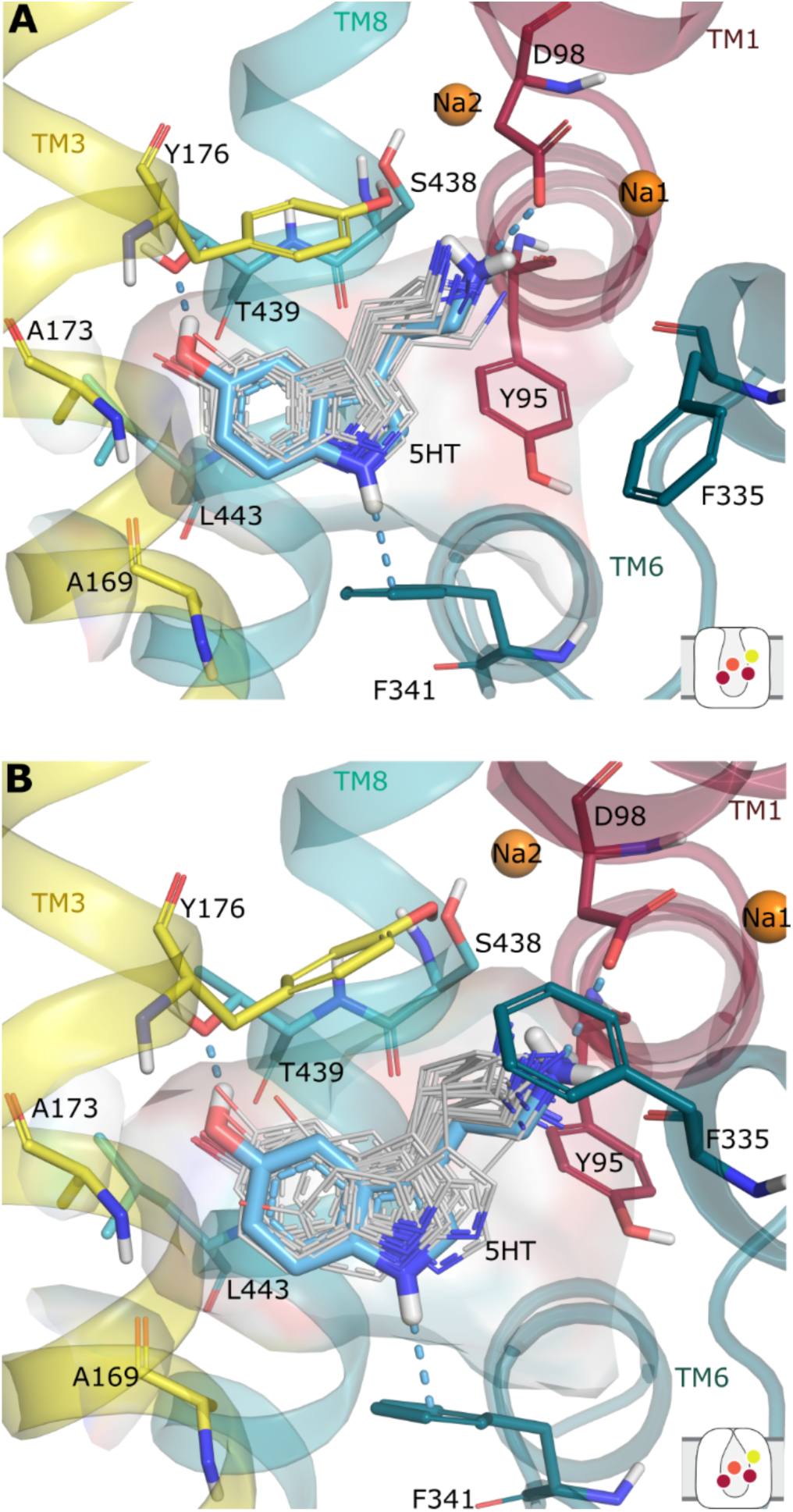
Conformations of 5HT in the most-favored clusters from induced-fit docking. Selected docking pose of 5HT (blue) within the cluster in A) the outward-open structure (PDB ID 5I71, 30 poses) and B) the outward-occluded model (57 poses). Previously-identified protein-ligand interactions with D98, F341, and T439 are observed in both cases (blue dashes).

### Binding pose of 5HT in the outward-open state

For the outward-open X-ray structure, the most-favored cluster of 5HT poses from the induced-fit docking was selected as that with the highest number of poses (30/74), as well as the best Glide and IFD scores on average (S10 Table). The protein-ligand interactions predicted by these poses were also consistent with previous studies. The top-ranked pose from this cluster, with a Glide Gscore of –9.1 kcal/mol, was used as a starting point for MD simulations (**Fig 5B**).

### Comparison of outward-occluded and outward-open states of hSERT

As mentioned in the Introduction, the outward-occluded conformation is, physiologically, the first state to be adopted after all ions and the substrate have bound. Unlike the outward-open state, the orthosteric binding site is covered and therefore inaccessible from both sides of the membrane [17]. The modeling approach applied here is based on our knowledge of other LeuT-fold transporters, and consequently the model displays similar characteristics (**Fig 6**). Specifically, TM1b and TM6a bend in to the extracellular vestibule towards TM3 and TM10, while EL4 functions as a lid and moves closer to EL6. However, the scaffold region and the lower half of the bundle helices remain unchanged.

**Fig 6:**
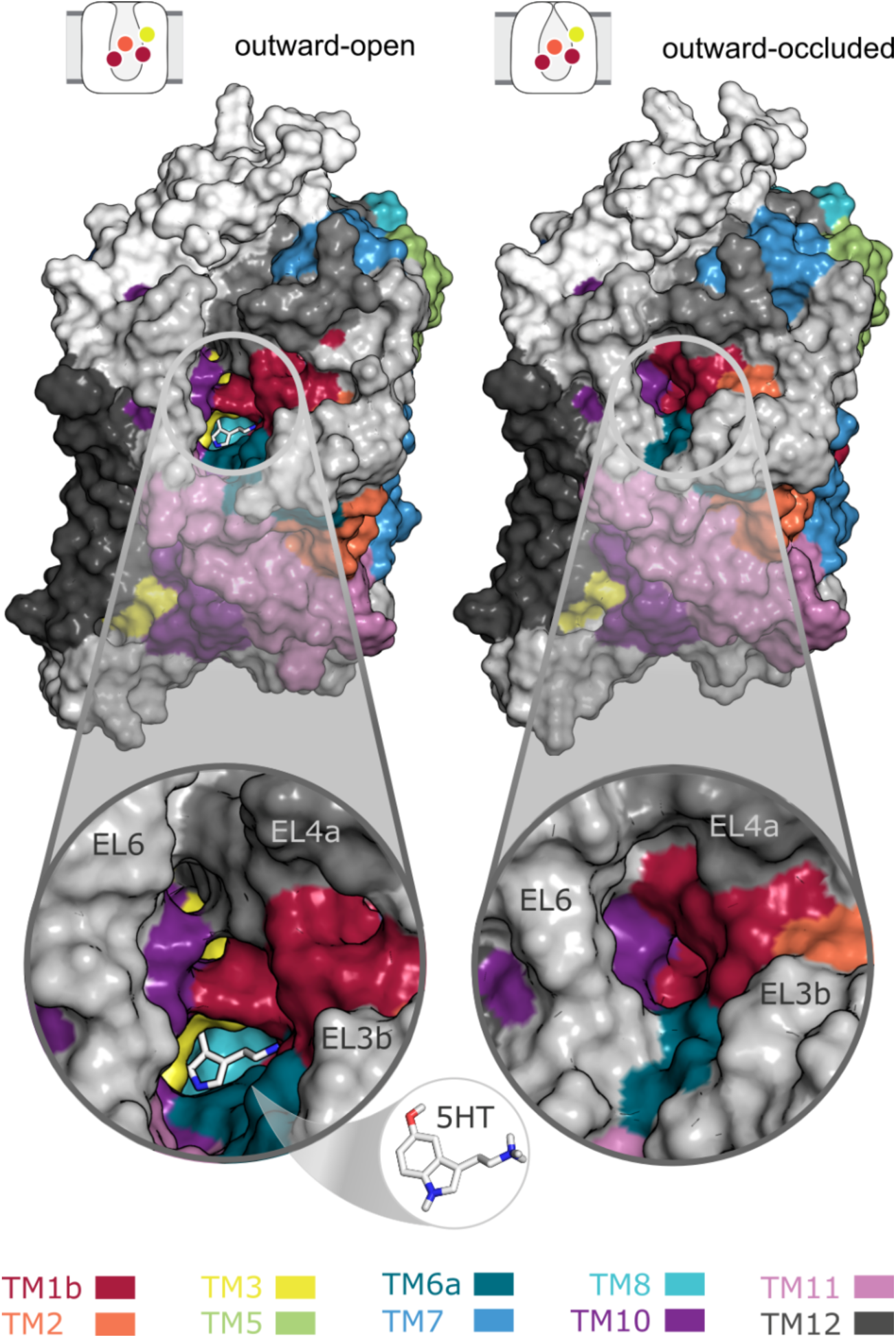
Differences between outward-open (left) and outward-occluded (right) structures of hSERT. The surface representation highlights the accessibility of the orthosteric binding site from the extracellular side. Specifically, the substrate molecule is visible from outside in the outward-open structure, but is covered in the outward-occluded structural model, where TM1b and TM6a bend over and close this same pathway from the synaptic cleft.

To explore the stability of the ion- and 5HT-bound outward-occluded model of hSERT and to gain insights into its behavior in the presence of thermal fluctuations and in an explicit lipid environment, we carried out multiple, microsecond-long MD simulations, and compared them with simulations of the outward-occluded complex. Both states proved to be stable over time: the average RMSD of the protein backbone (excluding the termini and EL2) in all simulations (8 μs) does not exceed 0.2 nm. However, the two protein conformations exhibited characteristic differences in their gating behaviors and protein-ligand interactions, as described below.

First, we tracked the interactions between residues forming the gates in the extracellular pathway **(Fig 7)**. In the outward-open state, R104 in TM1b and E493 in TM10 form transient interactions, although they do not close the pathway continuously. Thus, although the transporter is fully loaded and therefore ready for substrate translocation, the gates do not close completely, indicating that our simulations sample an energy minimum close to the outward-open state. In line with the behavior of the salt bridge, the hydrophobic lid formed by Y176 in TM3 and F335 in TM6a remains stably closed in simulations of the outward-occluded state, whereas this lid remains open in the simulations of the outward-open complexes. From these observations we conclude that our outward-occluded model remains in this conformation over time.

**Fig 7:**
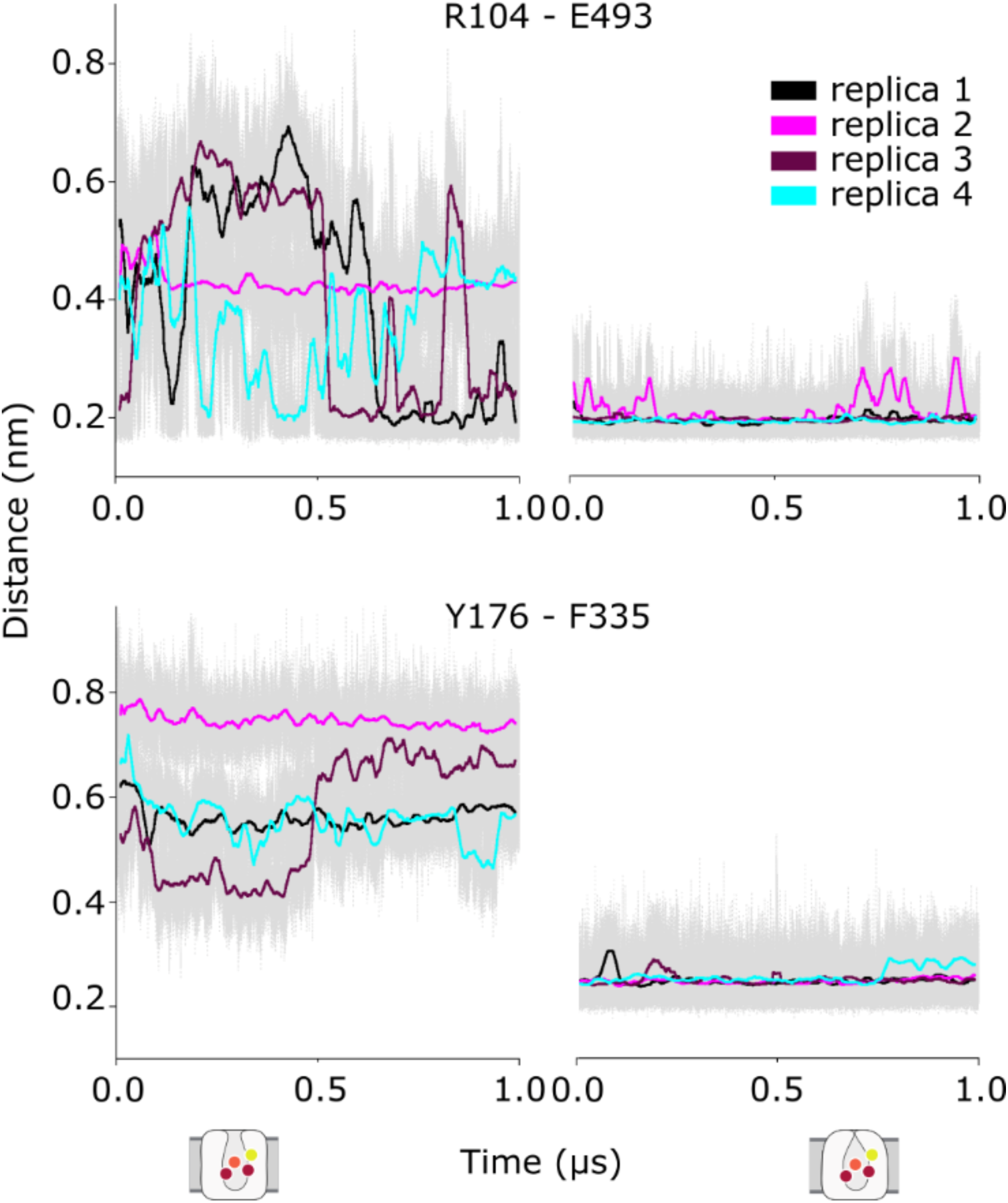
Behavior of the extracellular gates over time during molecular dynamics simulations of hSERT. The average minimum distances of the atoms between the R104-E493 and Y176-F335 gates are shown in 4 different trajectories for the outward-open (left) and outward-occluded states (right), with standard errors shown in the back (gray).

Although both conformations stay stable and 5HT remains bound, we were intrigued by the observation that the outward-open state does not drift towards the occluded state despite being occupied by all required substrates, and in contrast to previous reports for 5I6X [53]. We therefore investigated the protein-ligand interactions in the orthosteric binding site for potential explanations for the different interaction patterns in our systems. In both settings, the ligand stays stably bound, as reflected by the fact that the average RMSD of 5HT in all simulations (8 μs) does not exceed 0.1 nm. The positively-charged amine of 5HT interacts consistently with the negatively-charged side chain of D98. At pH 7, the 5HT primary amine is predicted to be protonated; the proximity with D98 likely increases this probability. The interaction with D98, however, cannot account for all the hydrogen-bonding potential for the primary amine, and thus we searched for additional interactions of the amine with residues in the vicinity. The backbone carbonyl oxygen atoms of Y95, F335, and S336 were all found to be in hydrogen-bonding distance. Monitoring these distances over time showed that all interactions could be sampled in both conformations, but stable contacts were only formed in the outward-occluded simulations **(Fig 8)**. These findings suggest that a stable network of Y95, D98, F335, and S336 coordinating the primary amine in 5HT may be required for the transition to the outward-occluded state.

**Fig 8:**
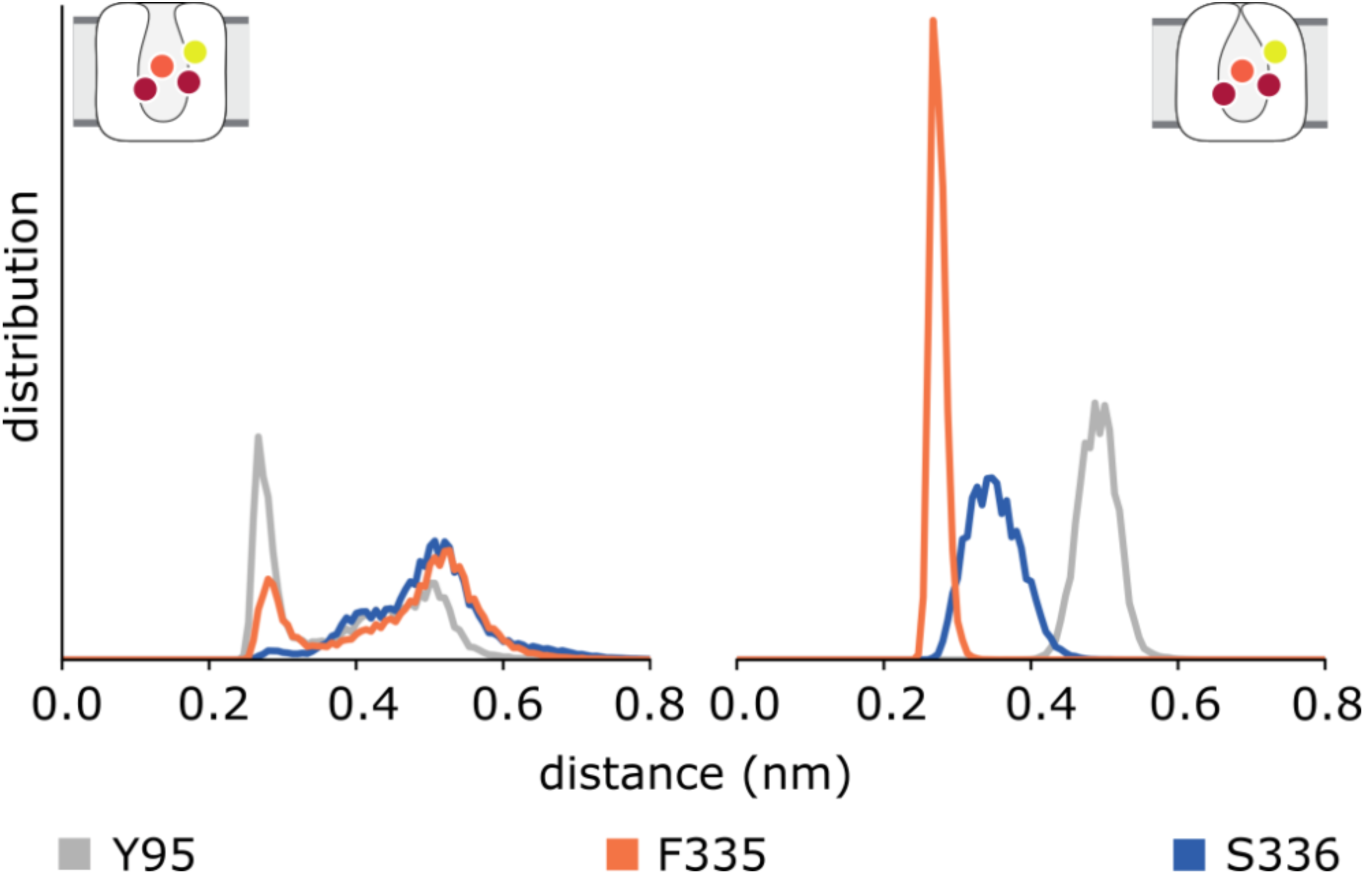
Interactions between the protonated nitrogen of 5HT and hSERT during molecular dynamics simulations. Aside from the salt-bridge with the deprotonated side chain of D98 (not shown), the amine coordinates the backbone oxygen atoms of Y95 (gray), F335 (orange), and S336 (blue). In the outward-open state (left), these distances increase over time, whereas they form a stable network in the outward-occluded state (right). Data are shown as normalized distributions over the combined simulation time.

### Putative allosteric consequences of 5HT binding

The orthosteric binding site is located in the center of the transporter between the extra- and intracellular pathways. Importantly, this site is formed in part by the conserved, unwound regions in TM1 and TM6, which enable the major conformational change between the different transporter states, at least in LeuT_Aa_ [27]. Given the timescale of our simulations, we do not expect substantial helical rearrangements in the intracellular half of the protein. Nevertheless, the observed differences in the orthosteric binding site have the potential to allosterically influence the intracellular interaction network [92–94]. We therefore investigated the distances between the residues forming the intracellular gate connecting the N-terminus, TM5, TM6, TM8, and TM9. Strikingly, the gates remain closed throughout the simulations of the outward-open conformation of hSERT, whereas they show increased conformational flexibility and dynamics in the simulations of the outward-occluded state **(Fig 9)**. Note that these motions do not indicate specific opening events of the intracellular pathway, e.g. the conserved residue pair W82-Y350 remains within a distance of 0.3 nm, but they do reveal a distinct behavior that appears to be induced by the occlusion of the pathway.

**Fig 9:**
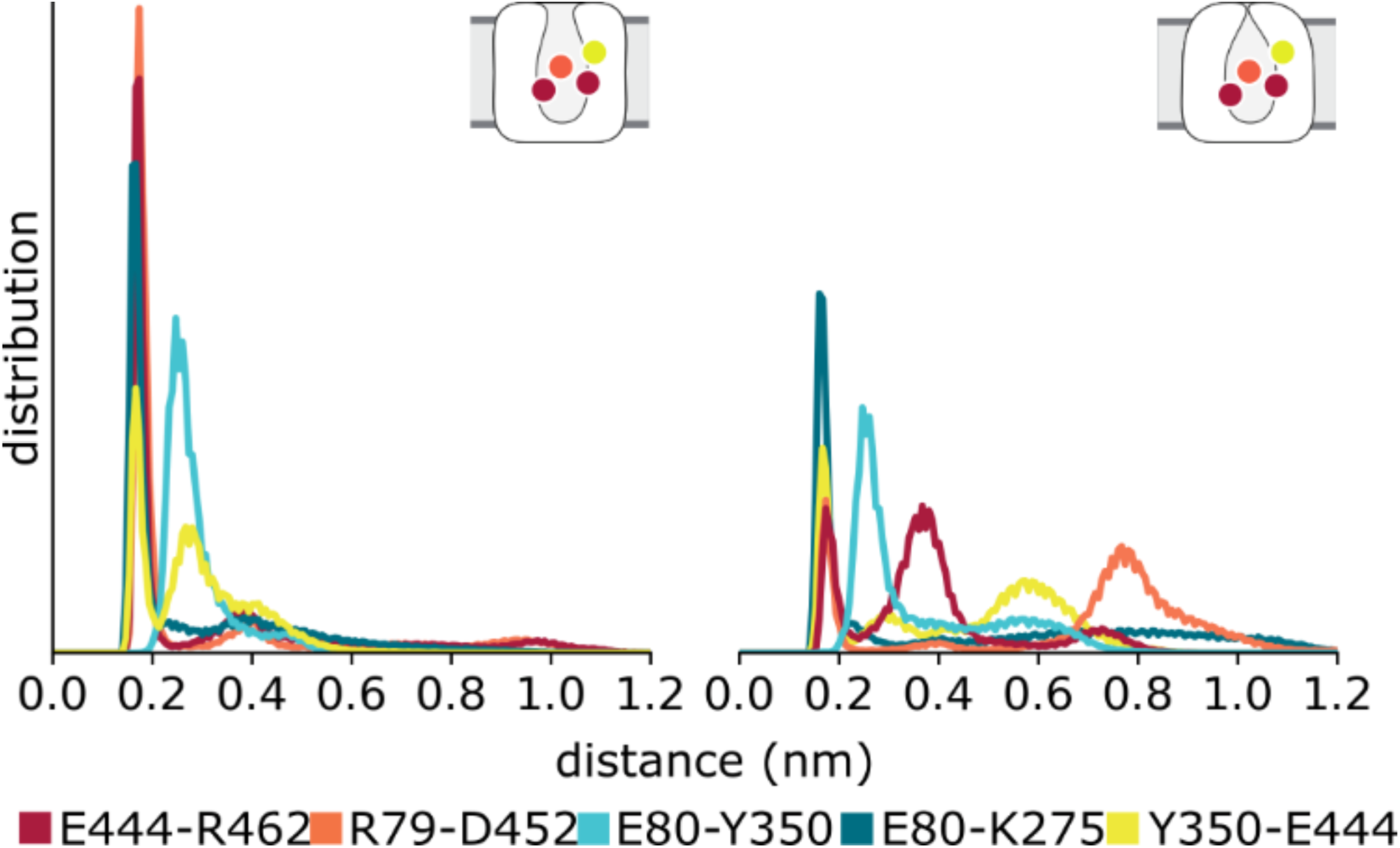
Distances between residues forming the intracellular gates during molecular dynamics simulations of hSERT. Residues in the cytoplasmic ends of TM5, TM6, TM8, TM9, and the N-terminus were tracked in simulations of the outward-open (left) and outward-occluded (right) states. The normalized distributions of distances between these gate-forming residues reveal a higher number of hydrogen-bond breaking events in the outward-occluded models.

Taken together, the distance analyses of the extra- and intracellular gates, and of the orthosteric binding site, predict subtle changes on the molecular level that may define the differences between the dynamics in the outward-open and outward-occluded states of hSERT.

### Hydration of the intracellular and extracellular pathways

The consequence of these observed differences can be quantified by the number of water molecules entering the transporter during the simulations **(Fig 10A)**. In the extracellular vestibule, we observe a clear disparity: in the outward-open state, a large number of (~10-20) water molecules occupies the pathway, whereas the closed gates in the outward-occluded state preclude hydration, allowing considerably fewer (~5) water molecules to occupy this region of the protein.

The hydration of the orthosteric site exhibits a pattern similar to that of the extracellular pathway. That is, more water molecules reach the core of the transporter when it is in the outward-open conformation, although the width of the distribution is smaller for both conformations. The latter observation presumably reflects the generally limited volume of the pocket and its occupation by 5HT, which provides a steric barrier to additional waters.

In the intracellular vestibule, in contrast, more water molecules enter in the simulations of the outward-occluded state than in those of the outward-open state. Although we do not observe any cases of ion or substrate unbinding to the cytoplasmic side, nor backbone conformational changes that might lead to such behavior, the influx of water in the simulations of the outward-occluded model may reflect a progression in the transport cycle due to allosteric consequences of substrate binding and extracellular pathway closure. Note, however, that none of these water molecules penetrate all the way to the substrate **(Fig 10B)**. In the simulations of the outward-open structure, the number of water molecules in the cytoplasmic side of the protein exhibits relatively large fluctuations both within and between trajectories (S11 Fig).

## Discussion

In this study, we have built a comparative model of hSERT in an outward-occluded state based on structural information primarily from the resolved hSERT X-ray structure itself. This was accomplished by fitting the upper bundle parts to the corresponding region in LeuT_Aa_ 3F48 before using them as a template combined with the remaining parts of the outward-open hSERT (5I71). We emphasized the model selection and refinement steps, and integrated knowledge from 5I71, 5I6X, and 3F48 to the greatest degree possible. The docked orientation of 5HT in its orthosteric binding site in both the outward-occluded model and the outward-open X-ray structure 5I71 were in excellent agreement with experimentally-validated induced-fit docking results obtained previously for a LeuT_Aa_-based model of hSERT [45]. Consequently, we conducted microsecond-long MD simulations of the ion- and substrate-bound complex, embedded in a membrane, to predict structural features that characterize the outward-occluded state. Simulations of the 5HT-bound outward-open structure of hSERT were used as a control.

Taken together, our data showed: (i) the extracellular gates remained closed; (ii) a stable interaction network coordinated the primary amine of 5HT in the orthosteric binding site; and (iii) fluctuations of the intracellular gates facilitated increased hydration of the cytoplasmic vestibule in the outward-occluded model.

Regarding the simulations of the outward-occluded model, it should be mentioned that the four replicas reported in this study were initiated from different frames of a single 2-μs trajectory (see Methods). Although this raises concerns that the trajectories are trapped in a local minimum, this decision in fact reflects our primary aim, namely to characterize the features of the outward-occluded state. Thus, we focused our efforts on trajectories in which the extracellular gates remained closed throughout the simulation, while trajectories in which the extracellular pathway stochastically reopened were discarded. To date, MD simulations of the major transition of hSERT from outward-open to inward-open conformations have not been reported. This is perhaps not unexpected, given that the time scale of a single out-to-in transition event in hSERT is ~10,000 μs [95–97], and thus, outside available classical MD simulations with current resources. Observations of intracellular pathway opening in hSERT, sufficient for sodium to diffuse away from its binding site, have been reported, albeit with the caveat that the starting point for the simulations was a LeuT_Aa_-based homology model [49]. However, in a recent study, Zeppelin et al. reported simulations of outward-open hSERT (using 5I6X) into which 5HT was placed according to Celik et al. [45]. In those simulations, the extracellular gate residues Y176-F335 also fluctuate between open and closed positions, in a similar way to the simulations of the outward-open state reported here (using 5I71) [53].

The inherent stochasticity of molecular dynamics simulations is also highlighted here by the unbinding of one of the bound chloride ions during one of the four trajectories of the outward-open state (S12 Fig). Although we did not observe rebinding during the length of the trajectories, such an unbinding event is not entirely unexpected, given the accessibility of the sites to the extracellular solution in that state.

It is noteworthy that all reported structures of hSERT lack the flexible C- and N-terminal ends of the protein, specifically, the regions comprising at least residues 1-73 and 616-630. The N-terminus has been implicated in amphetamine-induced efflux by hSERT [98], and in PIP2-mediated pathway opening in hDAT [99] and therefore the absence of residues N-terminal to position 74 might bias the intracellular gating behavior reported here. However, the fact that differences were observed in the transport-related elements of the protein for the different states suggests that the missing terminal regions are not essential for sodium-dependent serotonin uptake.

Very recently, Gouaux and coworkers reported cryo-EM structures of hSERT bound to ibogaine in outward-open (PDB code 6DZY chain A, hereafter referred to as 6DZY), outward-occluded (PDB code 6DZV chain A, hereafter referred to as 6DZY), and inward-open (PDB code 6DZZ chain A, hereafter referred to as 6DZZ) states. These structures are consistent with our assumption that the defining feature of the conformational change between the outward-open and outward-occluded states is the tilting of TM1b and TM6a to close the extracellular pathway [15]. After structural superposition of the backbone atoms in the hash domain, TM1b and TM6a of the outward-occluded model are found approximately mid-way between the equivalent helices in the outward-occluded and inward-open structures (Fig 11). As a quantitative measure of the extent of pathway closure, we measured the angle of these helices relative to the outward-open conformation (6DZY). Although TM6a is tilted to a similar extent in the outward-occluded model and the outward-occluded structure (6DZV) at ~8° ± 2° (mean and stdev over simulation time, calculated from steps at 20 ns intervals) and ~7°, respectively, TM1b is more tilted in our model (~17° ± 2°) than in the outward-occluded structure (6DZV; ~6°). These data suggest that ibogaine prevents TM1b from moving further into the pathway, but not TM6a. Importantly, this comparison also reveals that the helices are predicted to move in the same direction as the corresponding helices in the inward-facing structures of hSERT.

**Fig 10:**
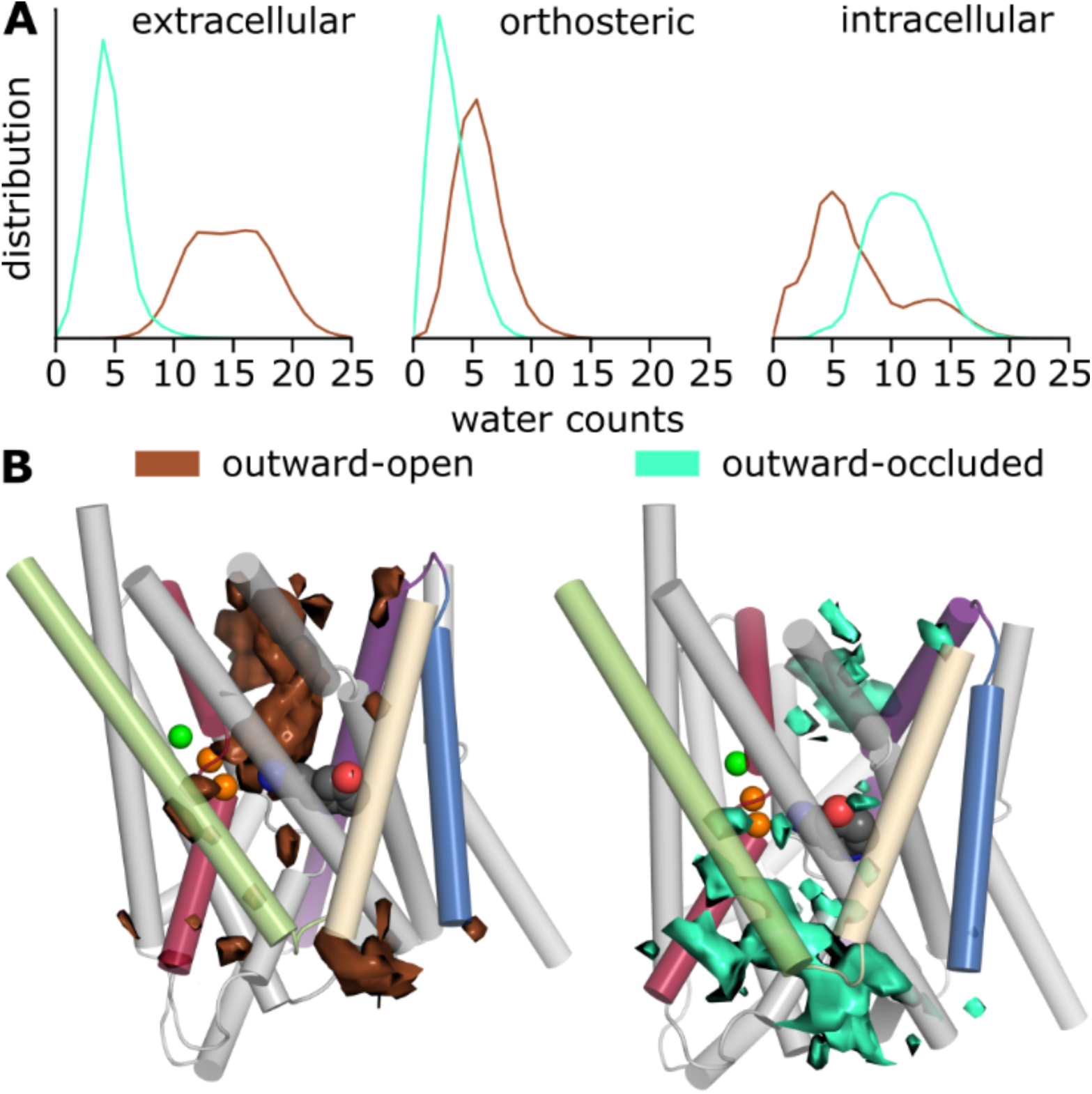
Hydration of the pathways in simulations of hSERT. A) The number of water molecules entering the central binding site, intra- and extracellular vestibules is shown as a normalized distribution over the total simulation time for the outward-open (brown) and outward-occluded states (green). B) Water occupancy within 20 Å of 5HT over the trajectory (1 µs simulation time) of one representative run for each state. The protein is shown as cylinders, highlighting specific helices colored according to Figure 2. The ligand (with C atoms in dark-gray) and sodium (orange) and chloride (green) ions are shown as spheres.

**Fig 11:**
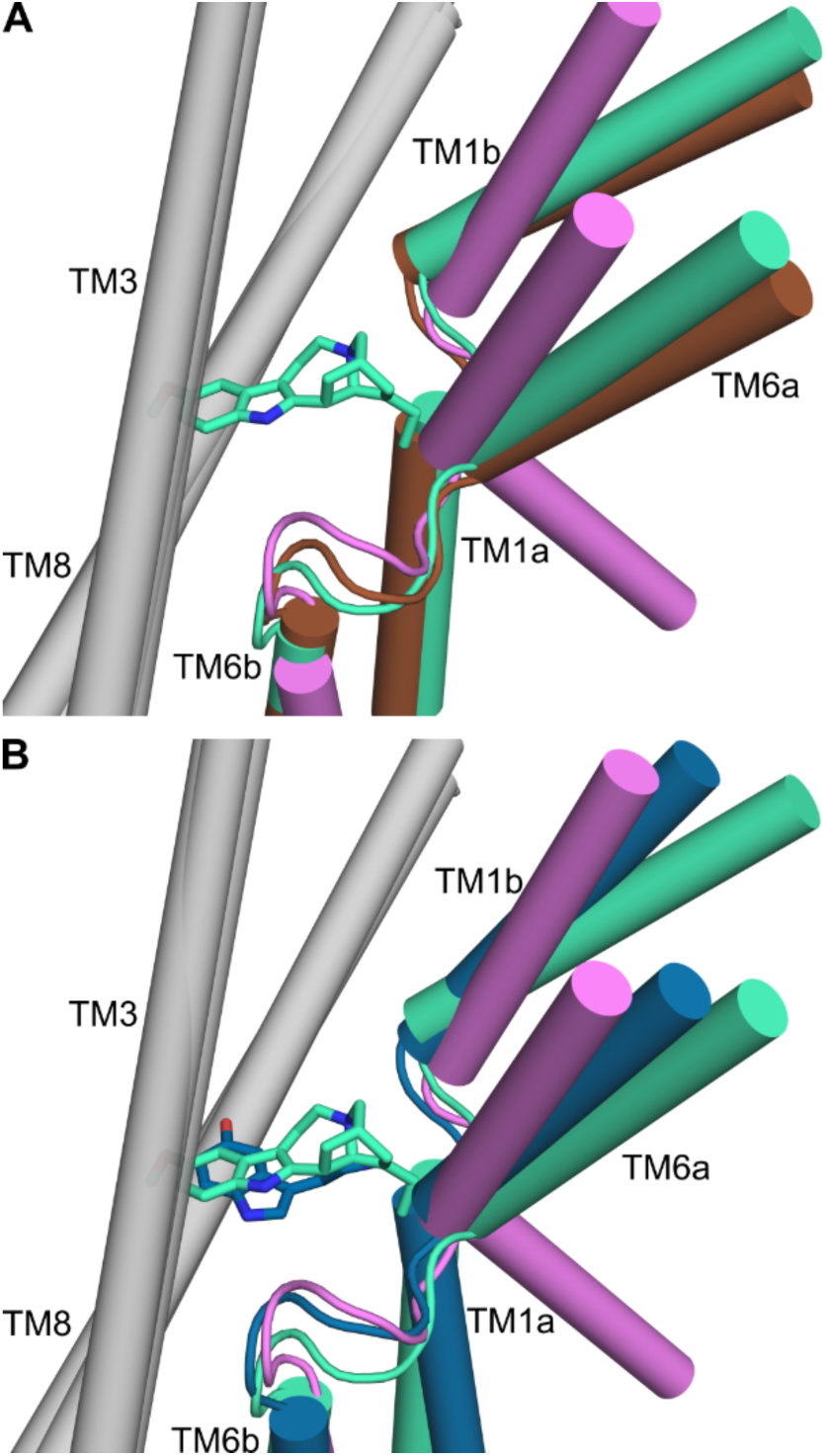
**Comparison of hSERT conformations:** shown are the ibogaine-bound cryo-EM structures in outward-open (brown), outward-occluded (green), and inward-open (pink) states, alongside the outward-occluded, 5HT-bound model (blue). TM1b and TM6a are shown in color, whereas TM3 and TM8 of the hash region are in grey. Ligands bound in the orthosteric binding site are ibogaine (green sticks) in outward-occluded 6DZV, and 5HT in the outward-occluded model (blue sticks). A) Comparison of the cryo-EM structures in outward-open (6DZY), outward-occluded (6DZV), and inward-open (6DZZ) states. B) The outward-occluded model presented in this study (blue, from the final frame of MD replica 2), bound to 5HT, has bundle helix positions midway between the outward-occluded and inward-open, ibogaine-bound cryo-EM structures. All structures in both panels A and B were aligned using the backbone atoms in TM3, TM4, TM8, and TM9, i.e. the hash domain.

Focusing on the extracellular gates, we also note that the side chain of F335 in TM6, which covers the orthosteric binding site from the extracellular pathway in LeuT, adopts a different orientation in our model than in outward-occluded structure 6DZY: the χ1 dihedral angle in the ibogaine-bound outward-occluded structure (–64°) is similar to that in the outward-open crystal structure (–70° in 5I71), whereas in our model the χ1 angle is −187° ± 12° (mean and stdev over simulation time), closer to that of the equivalent residue, F253, in the alanine-bound outward-occluded structure of LeuT_Aa_ (–166°).

Both of the aforementioned differences, namely the higher tilt and the different side-chain orientation, may reflect either the additional bulk of ibogaine (mol. wt. of 310.4 g/mol compared to 176.2 g/mol for 5HT) or differences in the mechanism of action of ibogaine relative to 5HT. The limited resolution of the cryo-EM structures should also be kept in mind. Finally, the binding mechanism of ibogaine is unknown, which leaves open the possibility that the recently-reported structures might not reflect true states in the substrate transport cycle.

Our conclusion that the conformation of the model is more occluded than the reported outward-occluded structure (6DZV) is supported by simulations of a 5HT-bound hSERT complex reported by Zeppelin et al., in which extracellular gate closure resulted in transient conformations in which TM1b is tilted to a similar extent as our model (~15° for the last frame of the trajectory). However, in their case, TM6a has not adopted the fully-occluded position, as it is only tilted by ~3° relative to the outward-open structure.

In conclusion, the predictions from our structural model are in line with the published computational and experimental data. Furthermore, they provide detailed hypotheses regarding the structural features characterizing the 5HT-bound, outward-occluded transporter, as distinct from the outward-open state. We anticipate that these predictions will aid in future studies elucidating the transport mechanism in hSERT.

## Acknowledgements

The computational results presented were obtained using the Vienna Scientific Cluster (VSC). We thank the National Heart Lung and Blood Lobos administrators for technical support, Richard Bradshaw for help with water analysis, Talia Zeppelin and Birgit Schiøtt (University of Aarhus) for sharing their simulation data, and Gary Rudnick for helpful discussions.

## Supporting Information

**S1 Table. Similarity of structural elements in hSERT and LeuT.** Results are reported as RMSDs (Å) and sequence identity (%) of the aligned segments. Note that the EL3TM6a segment was divided into three parts for the structure alignment.

**S2 File. Sequence alignment input for Modeller.**

**S3 File. Input file for CHARMM energy minimization.**

**S4 File. Ligand parameters for serotonin.**

**S5 Table. Ligand-protein distance restraints used for equilibration.**

**S6 Data. Final snapshots of four simulations of the outward-occluded model of hSERT.**

**S7 Data. Final snapshots of four simulations of the outward-open structure of hSERT.**

**S8 Tables. Structure quality scores and ion-protein relevant distances.**

**S9 Tables. Docking and clustering results for the hSERT outward-occluded model.**

**S10 Table. Docking and clustering data for the outward-open structure (5I71).** The most-populated clusters are defined as those containing >15% of all poses (i.e. >10 poses per cluster). The number of poses per cluster (or for the whole dataset) are reported alongside the mean ± standard deviations of the Glide Gscore and IFD scores for all poses in that cluster.

**S11 Fig. Water counts in the intracellular vestibule as a function of simulation time.** Number of waters in the intracellular vestibule during molecular dynamics simulations of hSERT, as a function of time. The plots show the underlying data for the outward-open (left) and outward-occluded (right) trajectories used to plot the distribution of the intracellular water counts in Figure 10A of the main manuscript.

**S12 Fig. Coordination chloride ions in simulations of the outward-open conformation.** Distance of the chloride ion from its initial position during the simulations of the outward-open model. The protein was least squares fitted to the initial frame of each of four replicas.

